# SARS-CoV-2 variants exhibit differential gut tropism and dysbiosis in Syrian golden hamsters

**DOI:** 10.1101/2023.06.19.545534

**Authors:** Santhosh K. Nagaraj, Christy M. Joy, Rishad Shiraz, Rohan Narayan, Sumandeep Kaur, Oyahida Khatun, Sagar Dubey, James Ng, Nischay Mishra, Shashank Tripathi

**Affiliations:** Emerging Viral Pathogens Laboratory, Infosys Wing, Centre for Infectious Disease Research, Indian Institute of Science; Bengaluru, 560012, India; Department of Microbiology and Cell Biology, Division of Biological Sciences, Indian Institute of Science; Bengaluru, 560012, India; Centre for Infection and Immunity, Mailman School of Public Health, Columbia University, New York, NY, USA

**Keywords:** SARS-CoV-2, VOCs, Syrian Golden Hamster, GI Infection, Gut Dysbiosis

## Abstract

Severe Acute Respiratory Syndrome Virus-2 (SARS-CoV-2) is a respiratory virus that uses angiotensin-converting enzyme 2 (ACE2) protein as an entry receptor. Thus, ACE2 expression levels in different tissues should dictate viral tropism. Notably, human ACE2 mRNA and protein levels are most abundant in the gastrointestinal tract, a pattern mirrored in Syrian golden hamsters. This prompted us to investigate the gastrointestinal tropism of SARS-CoV-2 variants of concern, including the Wuhan-like Hong Kong strain, highly pathogenic Delta, and highly transmissible but mildly pathogenic Omicron variants in hamsters. Delta was the most pathogenic in the respiratory and gastrointestinal tracts, followed by ancestral Wuhan-like and Omicron strains. In the gastrointestinal tract, viral RNA load was significant in the proximal organs such as the oesophagus and stomach, highest in the SI, and minimal to undetectable in the colon. Additionally, all three variants reduced fecal microbial diversity, with the Delta causing the highest decrease in observed features and phylogenetic diversity. Our findings highlight Delta’s stronger preference for the gastrointestinal tract, suggesting a link between high virulence and gastrointestinal tropism of SARS-CoV-2 variants of concern.

## INTRODUCTION

Severe Acute Respiratory Syndrome Virus-2 (SARS-CoV-2) emerged in Wuhan, China, in December 2019 ^1^, rapidly spread across the world, causing the COVID-19 pandemic that has taken more than 7 million human lives worldwide ^2^. It is an enveloped virus with a single-stranded positive-sense RNA genome ^3^. Its virions are studded with spike ^4^, which binds to the angiotensin-converting enzyme 2 (ACE2) receptor on the host cell to facilitate entry, that also requires cleavage of the spike by host proteases like transmembrane protease serine 2 (TMPRSS2) at the cell surface and cathepsin L in the endosomes ^5^. Its error-prone genome replication allows the accumulation of mutations ^6^, which led to the emergence of novel variants of concern (VOC), such as Alpha, Beta, Delta, and Omicron variants, with enhanced immune escape, transmission, and altered virulence ^7^. While vaccines and antivirals have helped to control the COVID-19 pandemic, SARS-CoV-2 continues to evolve in humans and wild animals ^8^.

COVID-19 is primarily a respiratory illness; however, in severe cases, extrapulmonary infection can appear, leading to gastrointestinal (GI), cardiovascular, hematologic, renal, neurologic, ophthalmologic, and dermatologic symptoms ^9,10^. The widespread expression of ACE2 across host tissues, along with patient co-morbidities and compromised immunity, contributes to the virus’s broad tropism and diverse disease symptoms ^10,11^. Several studies have highlighted GI symptoms in COVID-19 patients, especially in severe and long-COVID cases ^11^. Few studies have investigated SARS-CoV-2 infection in the GI tract and associated gut dysbiosis using animal models. However, strain-specific differences remain largely unexplored^12–14^. We examined the human proteome database to understand the tissue-specific expression of ACE2 and found its highest expression in the GI tract, especially the small intestine (SI). Syrian golden hamsters have been the preferred animal model for studying SARS-CoV-2 respiratory infection and disease and for testing the efficacy of vaccines and antivirals ^15^. Furthermore, we observed that the ACE2 expression pattern is recapitulated in the GI tract of hamsters, prompting us to use them to examine SARS-CoV-2 GI-tropism and pathology ^16,17^. Specifically, we compared the SARS-COV-2 Hong Kong strain, highly pathogenic Delta, and high transmissible Omicron VOCs ^18–22^ for their infection, replication, and pathology in the respiratory and GI tract of hamsters. Overall, we observed that Delta VOC replicated at a higher level but also showed higher inflammatory condition as seen with hematological parameters or cytokine levels and caused greater GI and respiratory pathology and a more significant disruption in the gut microbiome. This suggests that gut infection and dysbiosis are potential hallmarks of severe SARS-CoV-2 VOCs.

## RESULTS

### ACE2 and TMPRSS2 expression is higher in the GI tract compared to the respiratory tract of humans and hamsters

ACE2 is considered the primary receptor for SARS-CoV-2 entry and it is assisted by serine protease TMPRRS2. To understand the correlation between SARS-CoV-2 respiratory and GI tropism and the expression of ACE2 and TMPRSS2, we examined their mRNA and protein levels in human organs using the data available in the human protein atlas database ^36^. Interestingly, ACE2 mRNA and protein levels were significantly higher in the GI tract, particularly in the SI, compared to the lungs **(Figure 1A)**. A similar, though less pronounced trend was observed for TMPRSS2 at the protein level **(Figure 1B)**. We also assessed whether this expression pattern is recapitulated in uninfected hamsters by extracting mRNA from their lungs and GI organs. Like humans, hamsters showed the highest levels of ACE2 and TMPRSS2 in the SI **(Figure 1C and D).** This suggested that Syrian golden hamsters are suitable for studying the respiratory and GI tropism of SARS-CoV-2.

**Figure 1.**
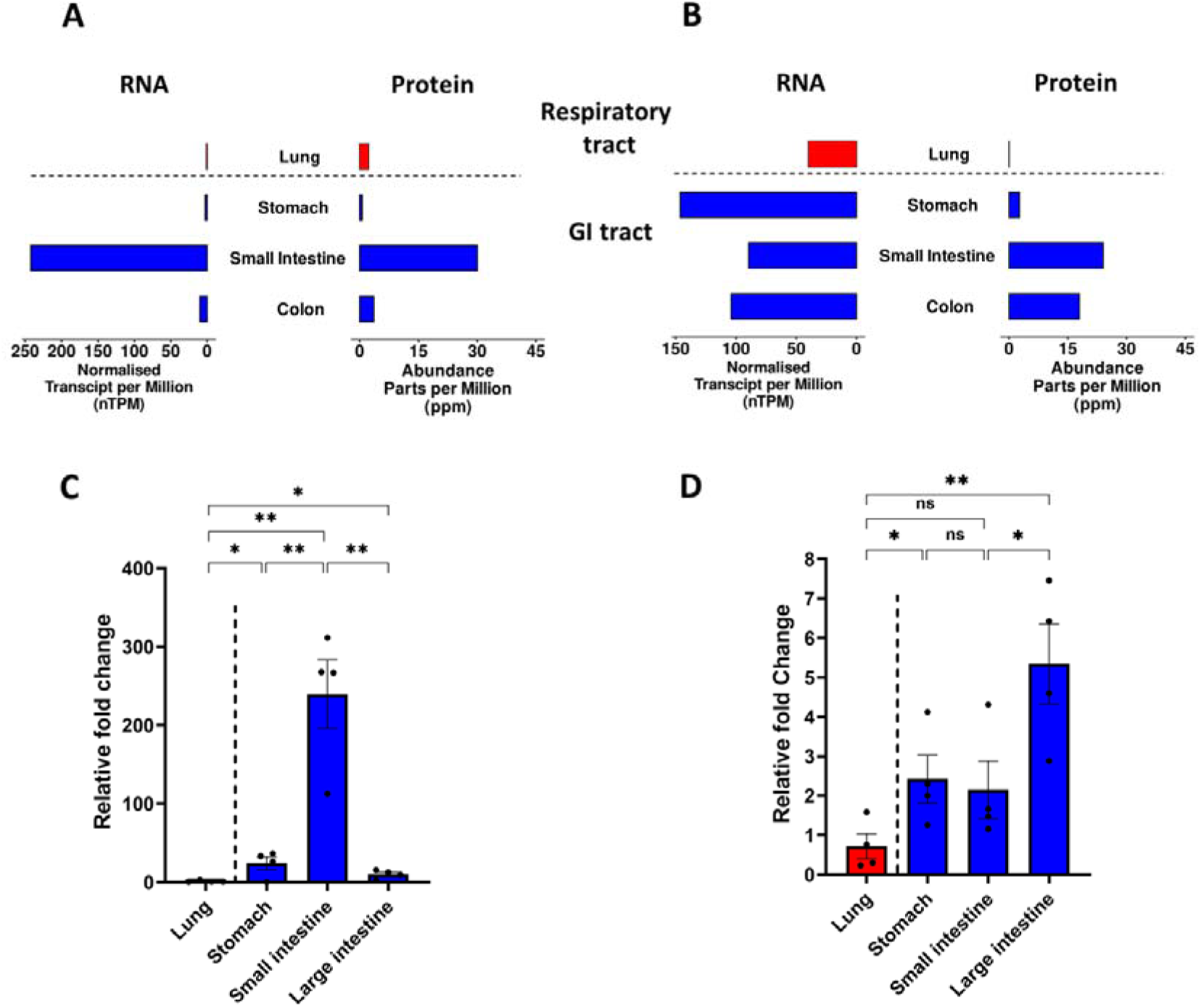
ACE2 and TMPRSS2 are expressed at higher levels in the GI tract of humans and hamsters. RNA and protein expression levels were obtained from the human protein atlas databases of **(A)** ACE2 and **(B)** TMPRSS2 in healthy human tissues such as the lung, stomach, SI, and colon. The RNA expression levels **(Relative to Lung)** of **(C)** ACE2 and **(D)** TMPRSS2 were measured in different tissues of hamsters (n=3 to 4), such as the lung, stomach, SI, and colon. *p < 0.05, **p < 0.01, ns - not significant. Error bars indicate Mean ± SEM.

### Comparison of clinical signs in hamsters infected with SARS-CoV-2 VOCs

For a comparative study of the disease caused by different SARS-CoV-2 VOCs, we chose Hong Kong, Delta, and Omicron variants. The comparison of their genome sequence revealed key amino acid differences with Omicron having most mutations particularly in the spike protein **(Figure 2A)**. To understand their differences in the pathogenicity and tissue tropism, hamsters were challenged with 10^5^ and 10^6^ PFU of each variant. The body weight, clinical signs, viral shedding in the nasal wash, viral replication and pathology in the lungs and GI tract, blood parameters, and fecal microbial diversity were assessed post-infection till 4 days post-infection (dpi) **(Figure 2B and C).** The Delta-infected hamsters had a maximum drop in body weight, followed by Hong Kong, and the least in the Omicron-infected animals **(Figure 2D).** The clinical signs also showed a similar pattern with comparable levels between Delta and Hong Kong variants with significantly milder and subclinical in Omicron infected animals (**Figure 2E).**

**Figure 2.**
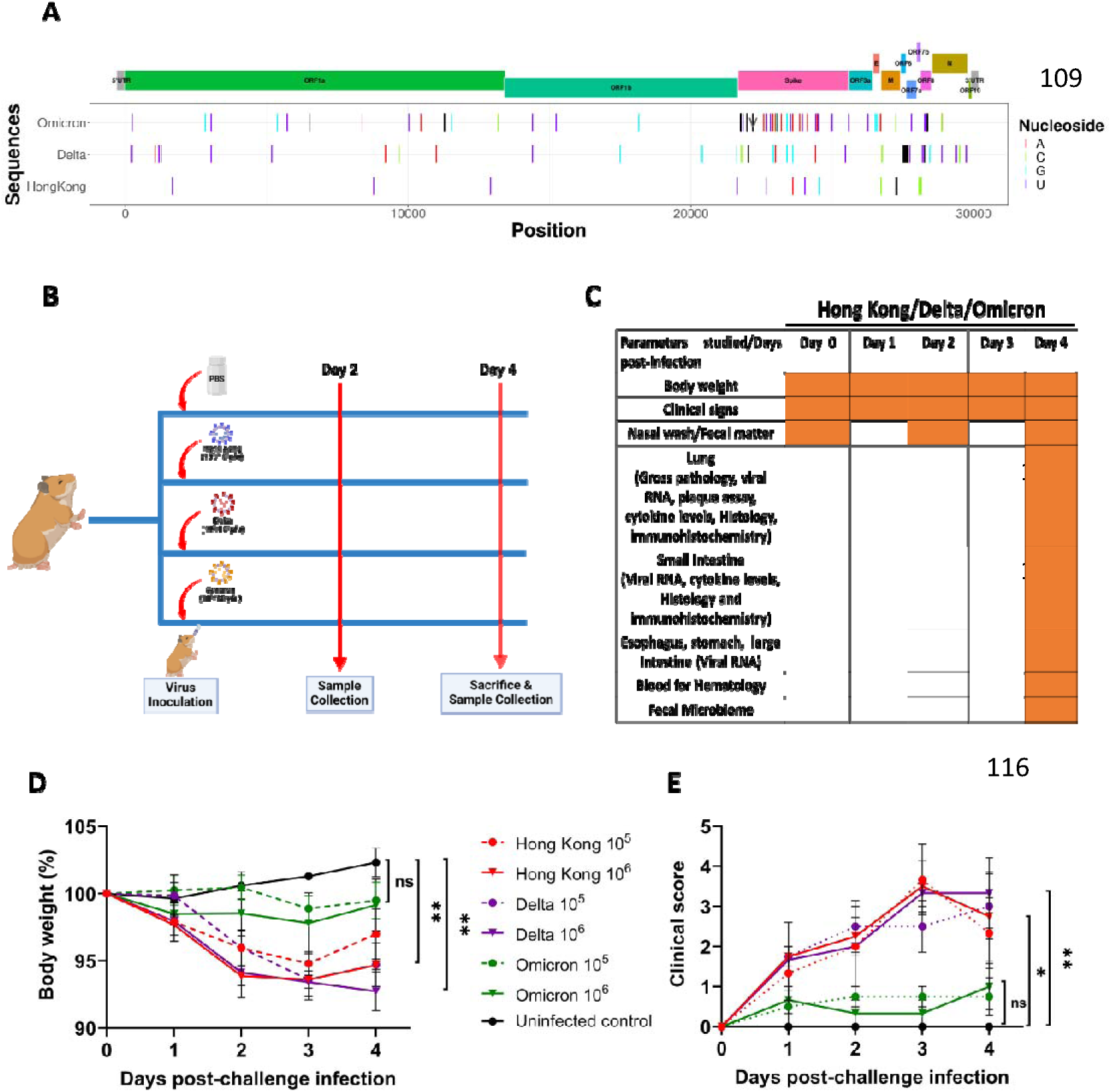
Genomic comparison of SARS-CoV-2 strains, hamster challenge study design and clinical signs in infected animals. **(A)** SARS-CoV-2 nucleotide mutation mapping of Hong Kong, Delta, and Omicron compared to the Wuhan-Hu-1 reference sequence. Single nucleotide polymorphisms and deleted nucleotide positions are marked using "|", with black indicating a nucleotide deletion and different colors denoting different nucleotide substitutions. Insertions are marked using "v" in dark grey color. The 5’ and 3’ untranslated regions are marked in grey, and the open reading frames (ORFs) and structural proteins are represented in different colors. **(B-C)** Outline of the experimental work plan followed in this study including the details on type of samples collected till the end of the experiment. **(D)** Percentage bodyweight of hamsters (n=3 to 4) monitored over 4 days post-infection (dpi), normalized to mean weight measured at day 0. **(E)** Clinical signs in hamsters including piloerection, abdominal respiration, hunched back, and lethargy was monitored over 4 dpi and scored from 1-4 based on severity. *p < 0.05, **p < 0.01, ns - not significant. Error bars indicate Mean ± SEM.

### Comparison of replication and pathogenesis in the respiratory tract of SARS-CoV-2 VOC-infected hamsters

We observed comparable viral RNA (vRNA) levels in the hamster’s nasal wash for all 3 VOCs on 2 dpi; but dropped significantly in Omicron-infected hamsters at 4 dpi **(Figure 3A and B).** However, in lungs, Delta-infected hamsters exhibited the highest vRNA load and infectious virus count, followed by Hong Kong and Omicron variants **(Figure 3C and D).** Immunohistochemistry (IHC) staining showed SARS-CoV-2 nucleocapsid protein in the lungs, with the highest levels in Delta, followed by the Hong Kong and Omicron groups **(Figure 3E and F).** Inflammatory responses, measured via lung/body weight ratio **(Supplemental Figure 1A)** and cytokine (IFN-LJ, IL-6 and TNF-α) mRNA levels **(Supplemental Figures 1B, C, and D)** were highest in Delta-infected hamsters, which also exhibited severe gross pathological changes than Hong Kong and Omicron variants at 4 dpi, consistent with RT-qPCR and plaque assay results **(Figure 4A and B)**. Further, at 10^5^ PFU dose, Omicron-infected hamsters showed no visible gross pathology.

**Figure 3.**
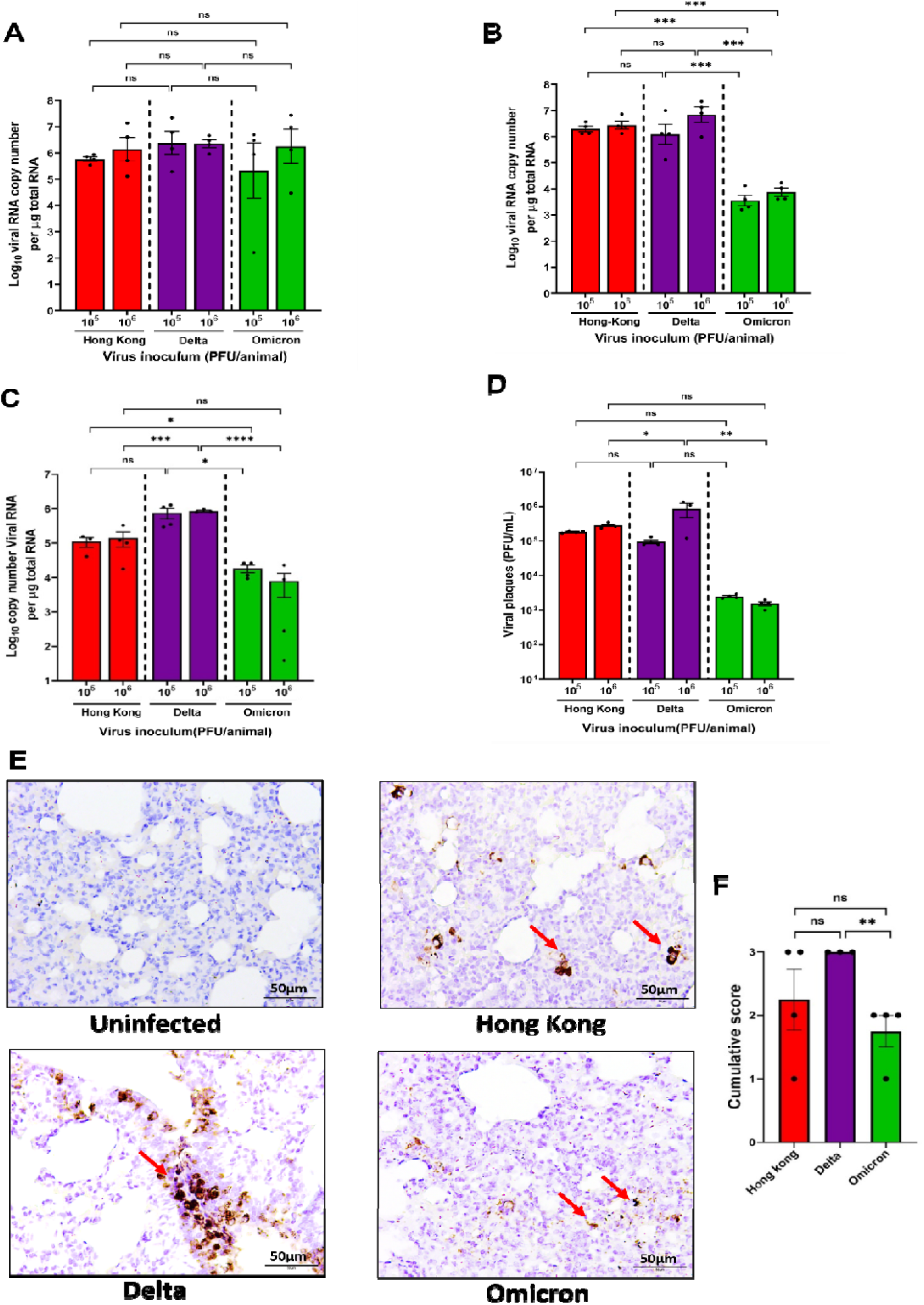
Viral load and nucleocapsid antigen levels in the respiratory tract of SARS-CoV-2 infected hamsters. Hamsters (n=3 to 4) were infected (10^5^ or 10^6^ PFU/100µL/animal) with Hong Kong, Delta or Omicron VOCs. Nasal washes were collected from animals at **(A)** 2 and **(B)** 4 dpi and viral RNA (vRNA) was estimated by RT-qPCR. **(C)** The total SARS-CoV-2 vRNA copy number in the lung was estimated by RT-qPCR. **(D)** Infectious SARS-CoV-2 titer was quantified by plaque assay. **(E)** Lung immunohistochemistry (IHC) staining (200x magnification) of uninfected and SARS-CoV-2 VOCs infected hamsters with the red arrow indicating the presence of SARS-CoV-2 N protein and **(F)** the cumulative scoring was also plotted. *p < 0.05, **p < 0.01, ***p < 0.01, ****p < 0.001, ns - not significant. Error bars indicate Mean ± SEM.

**Figure 4.**
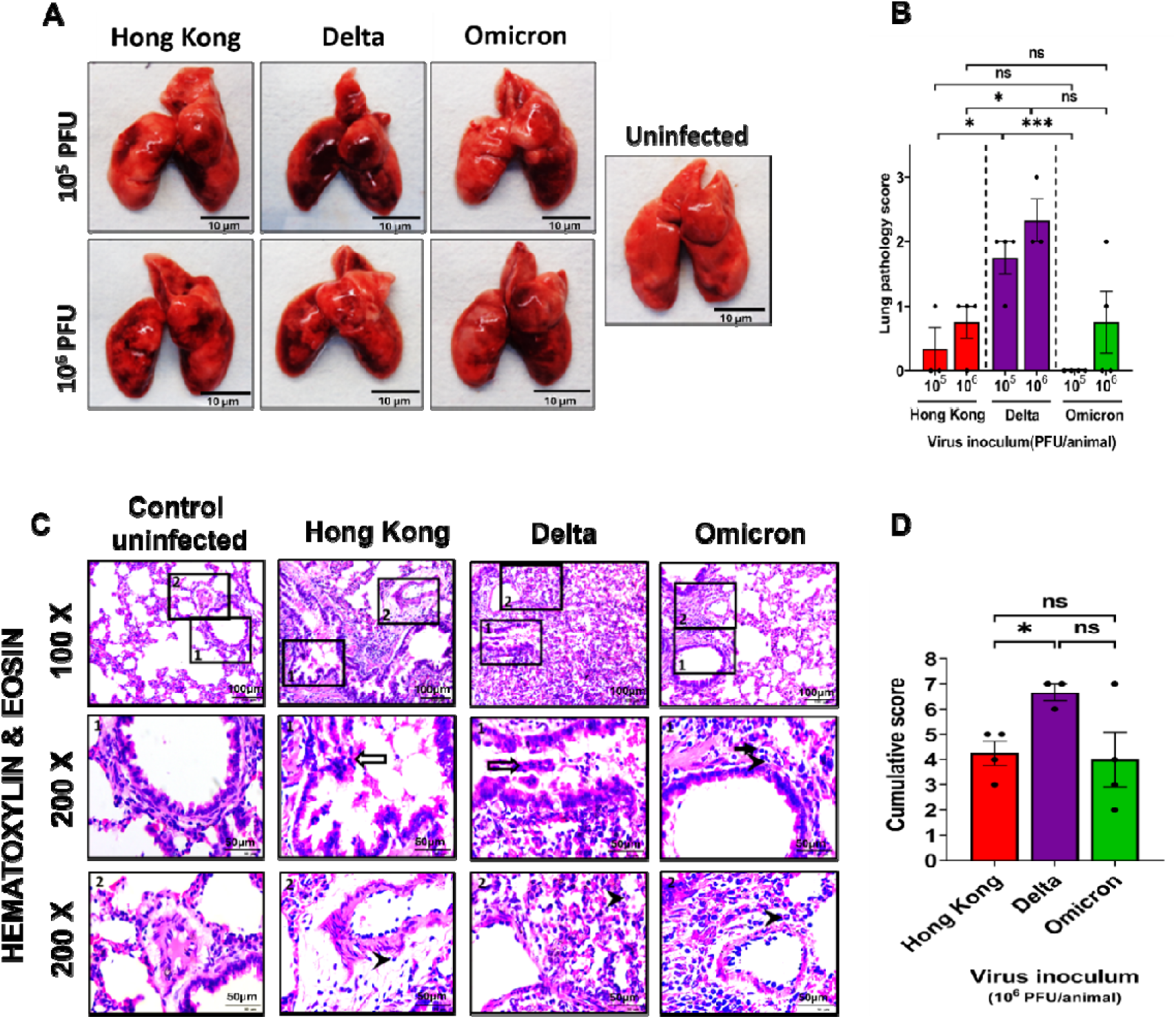
Lung gross and tissue pathology of SARS-CoV-2 VOC-infected hamsters. Hamsters (n=3 to 4) were infected (10^5^ or 10^6^ PFU/100µL/animal) with Hong Kong, Delta or Omicron VOCs and **(A)** lung images were captured at 4 dpi. **(B)** Lung gross pathology was scored 1-3 based on the severity of hyperemia. **(C)** Histopathology sections (10^6^ PFU) showed (1) bronchial epithelial necrosis/inflammatory changes (2) vascular inflammation (3) alveolar wall thickening and inflammatory cell infiltration. Bronchial and vascular changes are represented in insets 1 and 2 respectively (200x magnification), while alveolar thickening is in the first column of the figure set (100x magnification). White arrows indicate denuded necrotized bronchiolar epithelial cells admixed with inflammatory cells in the airway lumen, whereas inflammatory cell infiltration in peribronchiolar and perivascular areas are indicated as black arrows (lymphocytes) and black arrowheads (neutrophils). **(D)** Cumulative histopathology scoring for each group. *p < 0.05, **p < 0.01, ***p < 0.01, ns - not significant. Error bars indicate Mean ± SEM.

Histopathological features revealed broncho-interstitial pneumonia in Delta-infected hamster lungs, including vascular inflammation, alveolar infiltration, peribronchiolar infiltration with necrosis and mononuclear cell infiltrations in peribronchial and perivascular areas with occasional neutrophils. These lesions, mainly vascular inflammation, were more pronounced in Delta-infected hamsters, at 10^6^ PFU dose **(Figure 4C).** Cumulative lung histopathology scores (10^6^ PFU) were highest in Delta-infected hamsters, with significant differences compared to Hong Kong group **(Figure 4D)**. Omicron-induced lung pathology resembled that of Hong Kong infected hamsters.

Haematological analysis at 4 dpi (10^6^ PFU) revealed that Delta-infected hamsters had the lowest reticulocyte, basophil, and eosinophil counts but the highest WBC, neutrophils, and platelet counts **(Supplemental Figure 2A, B, and C)**. Further, Platelet distribution width levels were highest and lowest among Hong Kong and Omicron-infected hamsters, respectively **(Supplemental Figure 2C).**

### Comparison of viral replication in the GI tract of SARS-CoV-2 VOC-infected hamsters

Next, we examined the vRNA load in the esophagus, stomach, SI, and colon of hamsters infected with 10^5^ or 10^6^ PFU of SARS-CoV-2 VOCs. Omicron consistently showed the lowest RNA level across all these organs. Hong Kong and Delta-infected hamsters had similar RNA load in all organs, except in the SI, where Delta infection showed significantly higher vRNA levels **(Figure 5A and D).** Among GI organs, the vRNA was highest in the esophagus and lowest in the colon with the highest viral load in the SI of Delta-infected hamsters **(Figure 5E)**. Infectious virus (plaque assay) could only be detected in the SI of Delta-infected hamsters at 10^6^ PFU **(Figure 5F)**. IHC staining at 10^6^ PFU dose confirmed SARS-CoV-2 nucleocapsid protein in the SI of Delta-infected hamsters **(Figure 5G).**

**Figure 5.**
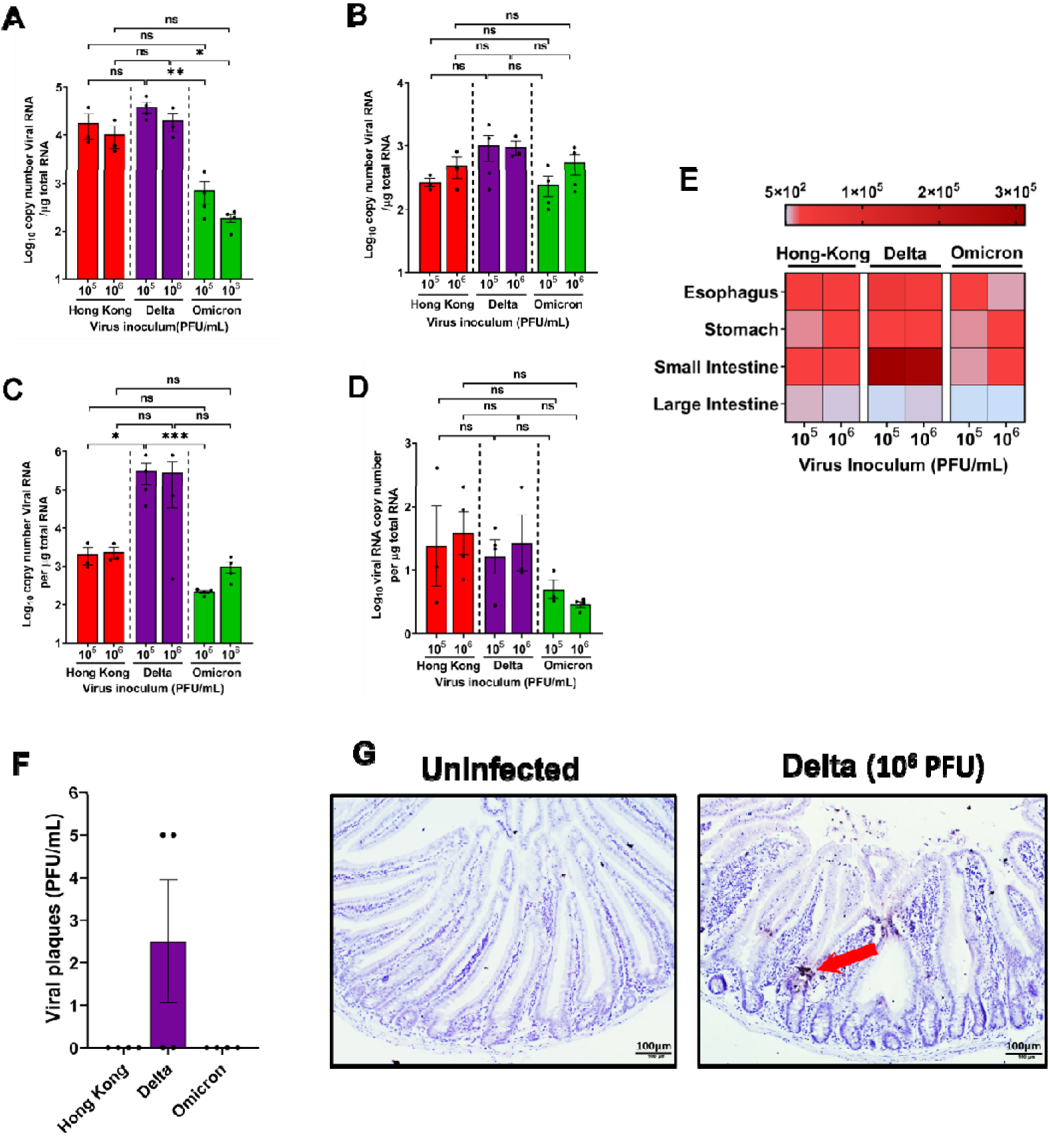
Viral load and nucleocapsid antigen levels in the GI organs of SARS-CoV-2 infected hamsters. Hamsters (n=3 to 4) were infected (10^5^ or 10^6^ PFU/100µL/animal) with Hong Kong, Delta or Omicron VOCs. Animals were sacrificed at 4 dpi and a portion of **(A)** esophagus, **(B)** stomach, **(C)** small, and **(D)** large intestines were stored In TriZol. Total RNA was extracted and vRNA copy number was quantified by RT-qPCR. **(E)** Heat map showing the measure of viral load compared among the virus variants and different organs tested in hamsters. **(F)** Viral plaque assay was carried out for the tissue homogenate of the SI. **(G)** IHC analysis with the red arrow indicating the presence of SARS-CoV-2 N protein. *p < 0.05, **p < 0.01, ***p < 0.01, ns - not significant. Error bars indicate Mean ± SEM.

### Comparison of inflammation and pathology in the SI of SARS-CoV-2 VOC-infected hamsters

To investigate SARS-CoV-2 VOC induced pathology in the SI, epithelial morphometric analysis was performed. An increase in crypt depth and reduction in villi/crypt length ratio is an indicator of tissue damage and pathology in the SI ^37^. The SI harvested from hamsters infected with the Delta variant had significantly increased crypt depth **(Figures 6A and B).** Although villi lengths were comparable across different VOCs, the villi/crypt length ratio was significantly reduced in the case of Delta-infected SI **(Figures 6A, C, and D)**. Overall, these data emphasize the SI mucosal damage and subsequent regenerative changes possibly caused by the Delta strain, as it also replicated to the highest level in the SI among different VOCs. Further, we assessed the inflammatory cytokines gene expression levels in the SI of infected small intestinal tissue. We observed that in the case of Delta-infected animals, interferon-gamma (IFN-LJ) **(Figure 6E)**, inteleukin-6 (IL-6) **(Figure 6F)**, and tumor necrosis factor (TNF-α) **(Figure 6G)** in the SI were relatively higher. Overall, Delta which replicates at a higher rate in the SI, also caused maximum inflammation and pathology in the SI of the hamsters, when compared to other SARS-CoV-2 variants.

**Figure 6.**
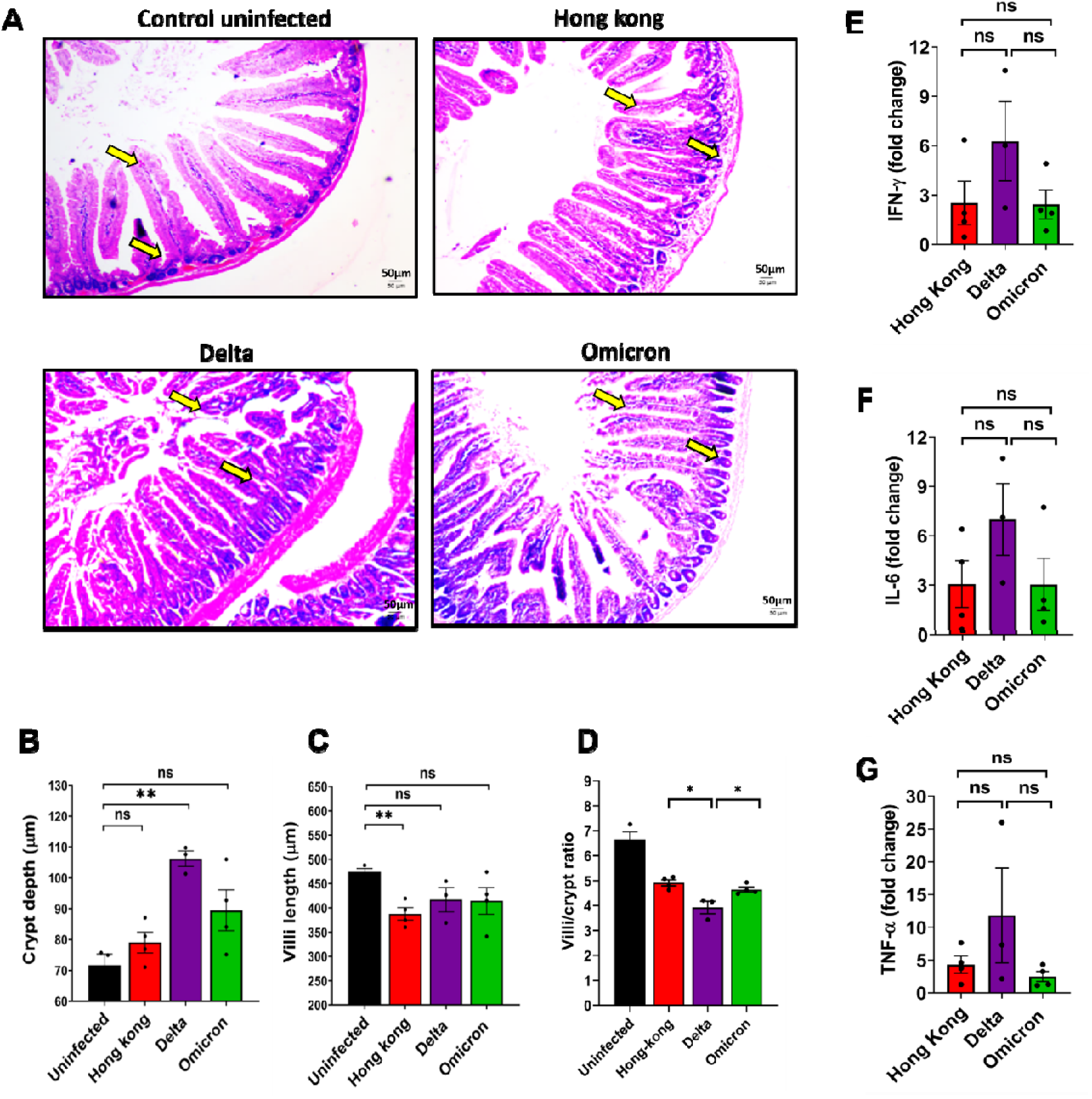
Tissue pathology and inflammatory cytokine in the SI of SARS-CoV-2 VOC-infected hamsters. Hamsters (n=3 to 4) were infected with 10^6^ PFU/100µL/animal of Hong Kong, Delta or Omicron VOCs. Animals were sacrificed at 4 dpi and SIs were collected and processed for histology. **(A)** Representative H & E-stained sections of duodenum of normal and different infected groups at 40x magnification showing the crypt depth and the villi (Yellow arrow). **(B)** Crypt depth and **(C)** villi length was quantified using ImageJ/Fiji. **(D)** Villi**/**crypt ratio was also measured. Data from 8-10 villi-crypt units per animal were measured. RNA was extracted from the SI and quantified the expression levels of IFN-D **(E)**, IL-6 **(F)**, and TNF-α **(G)** by RT-qPCR. *p < 0.05, **p < 0.01, ns - not significant. Error bars indicate Mean ± SEM.

### SARS-CoV-2 infection causes VOC-specific gut dysbiosis in hamsters

We further investigated the impact of VOC-specific variation in the gut microbiome by analysing fecal microbiota. All three variants showed reduced observed features (microbial richness) in fecal samples at 4 dpi, with Delta causing highest reduction, followed by Hong Kong and Omicron variants **(Figure 7A)**. Similarly, Faith’s phylogenetic diversity (Phylogenetic richness) showed greatest decline with the Delta variant, followed by Omicron and Hong Kong, compared to uninfected controls **(Figure 7B)**. We analyzed beta diversity to compare microbial compositions **(Figure 7C)**. The beta diversity (Jaccard distance) parameter showed distinct microbiota among all infected groups with the Delta displaying the most divergent microbial population compared to uninfected group.

**Figure 7.**
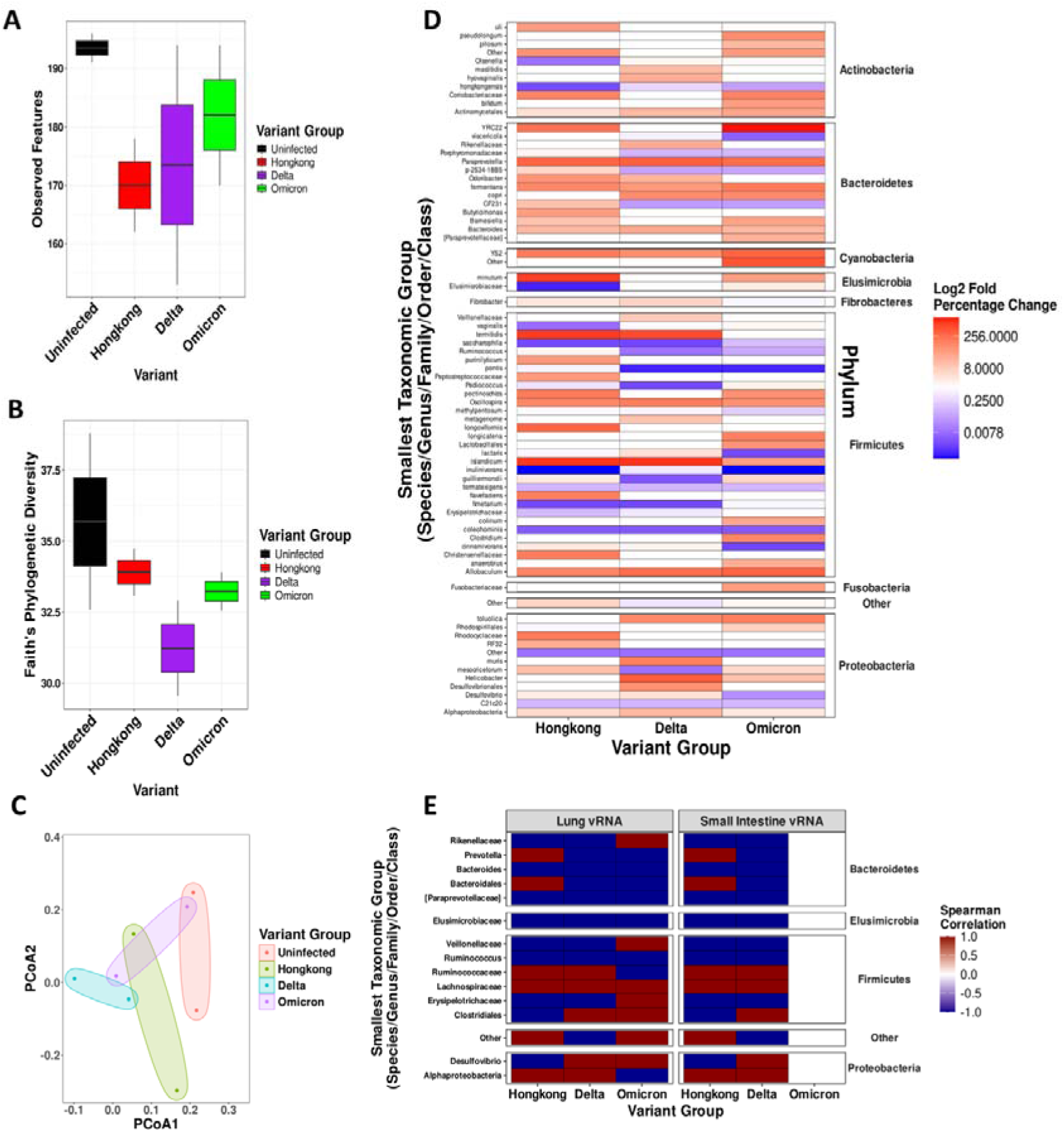
Higher gut dysbiosis in hamsters infected with Delta among the SARS-CoV-2 variants studied. Hamsters (n=3 to 4) were infected with 10^6^ PFU/100µL/animal of Hong Kong, Delta or Omicron VOCs. Fecal matter was collected on 4 dpi and microbiome levels were assessed by NGS sequencing. Alpha diversity was measured as **(A)** Observed features and **(B)** Faith’s phylogenetic diversity. **(C)** PCOA was plotted as a measure of Beta diversity. **(D)** Phylum-level microbial diversity was shown for different variants studied. **(E)** Spearman correlation between SARS-CoV-2 viral RNA (vRNA) and bacterial taxa were also assessed.

Comparative analysis of microbial proportions revealed specific changes following SARS-CoV-2 infection **(Figure 7D)**. While microbes such as *Clostridium islandicum*, *Paraprevotella*, *Allobaculum*, *Bacteroides*, *Oscillospira*, YS2 order bacteria, and *Macellibacteroides fermentans* were elevated over fivefold across all VOC infections, *Defluviitalea saccharophila* and *Lactobacillus coleohominis* showed consistent depletion. Variant-specific changes included elevations in *Lactonifactor longoviformis, Christensenellaceae, Rhodocyclaceae, Ruminococcus flavefaciens, Helicobacter mesocricetorum, CF231, Odoribacter, Clostridium purinilyticum, Olsenella uli, Peptostreptococcaceae,* and *Butyricimonas* (Hong Kong); *Helicobacter, Actinobacillus muris, Alphaproteobacteria, Desulfovibrionales, and Fibrobacter* (Delta); and *Dorea longicatena, Clostridium, Bifidobacterium pseudolongum, Lactobacillales, Rhodospirillales, Bifidobacterium bifidum, Fusobacteriaceae,* and *Oscillospira guilliermondii* (Omicron). Depletions unique to each variant were also observed, such as *Elusimicrobiaceae, Lactobacillus vaginalis, Oisenella,* and *Erysipelotrichaceae* (Hong Kong), *Pediococcus, Oscillospira guilliermondii*, and *Helicobacter mesocricetorum* (Delta), and *Papillibacter cinnamivorans, Ruminococcus lactaris, Barnesiella viscericola, Clostridium methylpentosum,* and *Desulfovibrio* (Omicron).

Spearman correlation analysis revealed associations between fecal microbiota and SARS-CoV-2 vRNA levels in the lungs and SI across variants **(Figure 7E**). While *Bacteroides*, *Paraprevotellaceae*, *Elusimicrobiaceae*, and *Ruminococcus* showed negative correlation, *Lachnospiraceae* showed a positive correlation across all VOCs. Variant-specific correlation patterns include *Prevotella* and *Bacteroidales* (positive with lung vRNA for Hong Kong but negative for Delta and Omicron) and *Rikenellaceae* and *Veillonellaceae* (positive for Omicron, negative for Hong Kong and Delta). *Ruminococcaceae* and *Alphaproteobacteria* correlated positively with Hong Kong and Delta vRNA but negatively with Omicron, while *Clostridiales* and *Desulfovibrio* showed opposite patterns (negative for Hong Kong and positive for Delta and Omicron).

## DISCUSSION

Coronaviruses cause respiratory and GI infections in mammals and birds ^38^. Also, SARS-CoV-2 infects diverse cell types and has been detected in multiple tissues of COVID-19 patients ^39,40^. The ACE2 receptor, a key determinant of SARS-CoV-2 tropism, is highly expressed in the GI tract, particularly the SI, with similar patterns observed for the entry co-factor TMPRSS2. Their highest co-expression occurs in the upper respiratory tract and intestinal epithelial cells, implicating that in addition to respiratory organs, the GI tract can also act as a site for viral replication and pathogenesis. Indeed, SARS-CoV-2 has been shown to infect and replicate in GI cell lines and intestinal organoids ^41^. A study reported GI symptoms in 59.7% of severe or hospitalized COVID-19 patients ^42,43^. Notably, the Delta variant replicates more efficiently in human colon epithelial cells than Omicron and other strains ^44^. These findings suggest that GI tropism and pathology are features of virulent SARS-CoV-2 variants, warranting further experimental validation. We observed that ACE2 and TMPRSS2 expression in hamsters closely resembles humans, with higher levels in the GI tract, particularly the SI, than in respiratory tissues, making them a suitable model for SARS-CoV-2 GI pathogenesis. We compared infection (via the natural route), replication, and pathology of Hong Kong, Delta, and Omicron variants in hamster respiratory and GI tracts. Additionally, we examined infection at two different doses to assess dose-wise variations in pathology among these variants. Delta caused the most severe disease, followed by Hong Kong, while Omicron infections were subclinical. Further, the increased lung cytokines, neutrophil/lymphocyte ratio or platelet levels in Delta-infected animals could suggest higher inflammatory condition which could be beyond the higher replication ability of the Delta virus including key adaptive mutations in Delta that enhance viral entry, replication, and immune evasion. Also, the presence of significantly higher viral load, followed by higher cytokine response, lead to more severe pathology in Delta infected hamsters compared to other variants studied. These findings align with published studies, with few unique strain VOC-specific observations in our study ^41,45–48^. In the GI tract, Delta showed the highest vRNA levels in the SI, followed by the oesophagus, stomach, and colon, while Hong Kong peaked in the oesophagus and Omicron showed significantly lower levels overall. This reflects variant-specific GI tropism, with the Delta infected SI being most affected, aligning with ACE2 expression patterns ^49^. Delta also induced higher inflammatory cytokines and SI epithelial damage, including villous blunting and regeneration in the duodenum, similar to COVID-19 patients ^50^. This suggests a strong adaptation of Delta for intestinal replication, leading to increased inflammation and mucosal damage (e.g., crypt depth increase, villi shortening). While our study found no vRNA in fecal matter at 4 dpi, delayed fecal vRNA shedding have been reported ^51^. GI infection and stool vRNA have been observed in fewer than 53% of cases and fecal-oral transmission is largely ruled out ^40,51–54^. Furthermore, studies on humans and hamsters link SARS-CoV-2 infection (parental strain) to microbiome dysbiosis in the lungs, SI, and fecal matter ^13,55^. Expanding on this, we compared GI microbiome changes across VOCs in hamsters. While all VOCs reduced fecal microbiome alpha diversity, Delta caused the greatest disruption, potentially contributing to its heightened pathogenicity in the GI and respiratory tracts. Taxonomic analysis revealed increased opportunistic bacteria (e.g., *Clostridium islandicum* and *Paraprevotella*) and depletion of protective commensals (e.g., *Lactobacillus coleohominis*), which may exacerbate disease severity caused by Delta variant. VOC-specific changes included enrichment of *Ruminococcus flavefaciens* with the Hong Kong variant and depletion of anti-inflammatory *Oscillospira guilliermondii* with Delta, that may further influenced pathology. Correlations between bacterial taxa and viral RNA highlighted protective roles for *Bacteroides* and *Ruminococcus* and pathogenic associations for *Lachnospiraceae*, underscoring the complex microbiota-virus interplay in disease severity. These correlations were consistent across the lungs and SI for Hong Kong and Delta variants and all VOCs have caused gut dysbiosis, with the Delta inducing the most significant disturbance. Further, association of gut-lung axis in exacerbating respiratory inflammation and disease severity has been reported earlier ^13,56^ . In our study, we speculate that the observed gut dysbiosis in Delta-infected hamsters, could suggest a potential role for the increased pro-inflammatory cytokines (IFN-LJ, IL-6, TNF-α) which might have induced a systemic inflammatory response that affects the lungs.

In conclusion, the Syrian hamster model is permissive to SARS-CoV-2 in the GI tract, with the Delta variant showing heightened tropism and pathology, particularly in the small intestine. While Delta and Hong Kong VOCs had similar GI viral burdens, Delta caused more severe symptoms and inflammation, which may have contributed to its enhanced transmission and severity in human populations. The findings support the role of the gut-lung axis in SARS-CoV-2 pathogenesis, particularly for the Delta variant, which caused the most severe GI inflammation, microbiome disruption, and respiratory pathology. Examining these effects in a long-COVID model and using post-Omicron VOCs will be interesting. Although the hamsters support SARS-CoV-2 replication in the GI tract, they do not display GI symptoms, which is a limitation ^57^. Nevertheless, this study warrants further exploration of host factors required for GI tropism and associated disease by SARS-CoV-2 and other human coronaviruses.

## MATERIALS AND METHODS

### Virus stocks

SARS-CoV-2 Isolate Hong Kong (hCov-19/Hong Kong/VM20001061/2020, Cat No.: NR-52282), Delta variant (USA/PHC658/2021, lineage B.1.617.2; Cat No.: NR-55611) and Omicron variant (USA/PHC658/2021, lineage B.1.1.529; Cat No.: NR-56461) were obtained from BEI Resources, NIAID, NIH. All viruses were propagated and titrated by plaque assay in VERO-E6 cells, as previously described ^18^.

### Animal Handling

Male and Female Syrian Golden Hamsters (*Mesocricetus auratus*) of 6-7 weeks old (60-70g) were housed in the viral biosafety level 3 laboratory at, the Indian Institute of Science, Bangalore. Animals were maintained in individually ventilated cages at 23±1°C temperature and 50±10% relative humidity with 12h day/night light cycle with food and water *ad libitum*. An overdose of Ketamine (Bharat Parenterals Limited) and Xylazine (Indian Immunologicals Ltd) was used to sacrifice animals upon completion of the experiment.

### Animal grouping and virus infection

A total of 28 animals, divided into 7 groups with 2 males and 2 females per group **(Figure 2C)**, were intraperitoneally anesthetized (Ketamine-150mg/kg and xylazine-10mg/kg) and intranasally infected with either Hong Kong, Delta, or Omicron virus at the dose of 10^5^ or 10^6^ Plaque Forming Units (PFU)/animal in 100µL PBS (50µL/nostril). Fresh feces (TRIzol) and nasal washes (100µL PBS) were collected pre-infection, on 2 and 4 dpi. Animals were sacrificed on 4 dpi and blood was collected by cardiac puncture and stored at -80°C. Portions of organs (lungs, esophagus, stomach, SI, and large intestine) were stored in TRIzol (RT-qPCR) or DMEM (plaque assay) or 10% buffered formalin (histopathological analysis).

### Clinical signs and lung gross pathology

Hamsters were daily observed for the following clinical signs till 4 dpi and scored based on severity for lethargy/piloerection (none=0, mild=1, moderate=2, severe=3); abdominal respiration/hunched back (absent=0, present=1) and bodyweight loss (1-5%= 1; 6-10%=2; 11-15%=3). The lungs were scored for focal and diffused hyperemia (mild=1, moderate=2, severe=3). The total lung weight was recorded and images were captured. The lung-body weight ratio was calculated.

### Hematological analysis

The blood sample was collected from hamsters on 4 dpi to analyze the complete blood parameters using a VetScan analyzer (VSpro, ABAXIS).

### Plaque assay

The plaque assay was performed in 12-well plates to determine the viral titer in tissue homogenates. A 100µL of 10-fold diluted samples (DMEM with 2% FBS) was added in duplicates over VERO-E6 cells, incubated for 1h at 37°C. After inoculum removal, 1mL of 0.6 % Avicel overlay was added and incubated at 37°C with 5% CO2 for 48h. The overlay was removed post-incubation and cells were fixed with 4% formaldehyde in PBS and stained with 1% crystal violet to visualize plaques.

### RNA extraction and Quantitative Real-Time PCR (RT-qPCR)

Organs were processed using a FastPrep-24™ stainless-steel bead homogenizer (MP Biomedicals, USA), and total RNA was extracted using TRIzol (15596018, Thermo Fisher) as per the manufacturer’s instructions. A 10µL reaction mixture (100ng RNA/sample) in a 384-well block was used to quantify vRNA using AgPath-ID™ One-Step RT-qPCR kit (AM1005, Applied Biosystems). The comparative threshold (Ct) values were used to determine viral copy numbers via a standard curve generated with SARS-CoV-2 genomic RNA. The host cytokines mRNA (IFN-γ, IL-6, and TNF-α) levels were also quantified by preparing cDNA from 1µg RNA using PrimeScript^TM^ RT reagent Kit with gDNA Eraser (TaKaRa). RT-qPCR was performed using 2.5μL of cDNA and primer mix each and 5μL of SYBR™ Green PCR Master Mix (Applied Biosystems). The Ct values were presented as mean Log_2_ fold change compared to 18S RNA by the delta-delta Ct method. The primer and probe sequences are provided in **supplementary table 1.**

### Histopathology and Immunohistochemistry

Tissue specimens were fixed (10% buffered formalin), paraffin embedded, sections (4-6μm) were stained with Haematoxylin and Eosin (H&E) and examined under light microscope. Parameters such as alveolar infiltration/exudation, vasculature inflammation, and peribronchiolar infiltration with epithelial desquamation (lungs) and the crypt depth/villi length (SI) were measured and scored on a scale of 1 to 3. Similar sections were used for quantifying the SARS-CoV-2 nucleocapsid protein (Novus Biological, NB100-56576) by IHC as previously described ^23^. Images were captured under a microscope.

### Microbiome analysis

Fecal DNA was extracted using QIAamp PowerFecal Pro DNA Kit (Cat No. 51804) and 20ng/μl per sample was used for 16S-EZ library preparation, targeting the V3-V4 regions with the MetaVX 16S rDNA Library Preparation Kit (Genewiz, NJ, USA). Sequencing was performed on an Illumina MiSeq platform (2×250 bp paired-end) and base calling with built-in Illumina software. Base FASTQ conversion were carried out using bcl2fastq 2.17 software. Quality-filtered reads were processed in Qiime2 (v2024.2.0), where paired-end reads were merged, and barcodes and primers were removed ^24^. Demultiplexing and denoising were performed with the DADA2 plugin to eliminate poor-quality, homopolymeric, and chimeric sequences ^25^. Samples were rarefied at 5647 reads to normalize sequencing depth, and amplicon sequence variants (ASVs) were assigned taxonomies using the Greengenes database ^26^. Alpha diversity (Observed features, Faith’s phylogenetic diversity) and beta diversity (Jaccard Distance) metrics were computed with Qiime2 ^24,27,28^. Relative changes in operational taxonomic units (OTUs) were assessed against the uninfected group to evaluate microbial shifts. Spearman correlation was used to examine the association between SARS-CoV-2 vRNA levels (lung and small intestine) and microbiome OTUs. Data visualization and statistical analyses were performed in RStudio using packages such as Tidyverse, gridExtra, RColorBrewer, purrr, qiime2R, ggsignif, ggpubr, rstatix, and upSetR^25,29–35^.

### Ethics statement

The study was approved and conducted in compliance with institutional biosafety (IBSC/IISc/ST/18/2021) and animal ethics (Ref: IAEC/IISc/ST/784/2020) guidelines.

### Statistical analysis

The data were analyzed using GraphPad Prism v 8.4.3 and represented as mean ±SEM. Statistical variations were determined by one-way ANOVA (Lung/bodyweight ratio, tissue vRNA load, cytokine mRNA level, ACE2/TMPRSS2 level, nasal wash and lung) or two-way ANOVA (Body weight, clinical signs) with Tukey’s multiple comparisons tests or by paired T-test (Lung histopathology). Values were significant when *p < 0.05, **p < 0.01 or ***p < 0.001 otherwise ns - not significant.

## AUTHOR CONTRIBUTIONS

Study Design: ST, NM; Resources: ST, NM; Experimentation and Data analysis: SK, CMJ, SN, RS, SK, OK, SD, JN; Manuscript Writing: ST, SN, RS, RN. All authors have given approval to the final version of the manuscript. ‡ Santhosh K. Nagaraj and Christy M. Joy contributed equally.

## Supporting information

Suuplemetary material

## ACKNOWLEDGMENTS

This work was supported by the DBT-BIRAC (BT/CS0007/CS/02/20) grant to ST. We acknowledge Viral BSL3 facility supported by BIRAC and Crypto-Relief. Microbiome study was supported by Yusuf Hameid Fellowship (Columbia University, NY) to NM.

## CONFLICTS OF INTERESTS

The authors have declared that no conflict of interest exist.

## DATA AVAILABILITY STATEMENT

The supplementary file contains the list of primers used for RT-qPCR, lung-to-body weight ratio, lung inflammatory cytokine levels, and haematological parameters studies in hamsters. All raw data related to the experiments shown in the figures are provided in the excel file.

## Notes

### Competing Interest Statement

The authors have declared no competing interest.

### Summary of Updates

We have added new data on Gut microbiome alteration induced by SARS-CoV-2 infection in Syrian Hamsters. Data show that SARS-CoV-2 VOC associated with severe diseases such as Delta infect the small intestine and induce greater gut dysbiosis, when compared to less pathogenic VOCs.

## REFERENCES

1. Hui DS, Azhar EI, Madani TA, et al. The continuing 2019-nCoV epidemic threat of novel coronaviruses to global health—The latest 2019 novel coronavirus outbreak in Wuhan, China. 2020;91:264–266.

2. (WHO) WHO. WHO Coronavirus (COVID-19) Dashboard. Geneva: WHO. . [cited 2024 Dec 22] 2024;

3. Gorbalenya AE, Baker SC, Baric RS, et al. The species Severe acute respiratory syndrome-related coronavirus: classifying 2019-nCoV and naming it SARS-CoV-2. 2020;5(4):536–544.

4. V’kovski P, Kratzel A, Steiner S, Stalder H, Thiel V. Coronavirus biology and replication: implications for SARS-CoV-2. Nature Reviews Microbiology. 2021/03/01 2021;19(3):155-170. doi:10.1038/s41579-020-00468-6

5. Koch J, Uckeley ZM, Doldan P, Stanifer M, Boulant S, Lozach PY. TMPRSS2 expression dictates the entry route used by SARS-CoV-2 to infect host cells. Embo j. Aug 16 2021;40(16):e107821. doi:10.15252/embj.2021107821

6. Phan T. Genetic diversity and evolution of SARS-CoV-2. Infection, genetics and evolution : journal of molecular epidemiology and evolutionary genetics in infectious diseases. Jul 2020;81:104260. doi:10.1016/j.meegid.2020.104260

7. Carabelli AM, Peacock TP, Thorne LG, et al. SARS-CoV-2 variant biology: immune escape, transmission and fitness. Nature Reviews Microbiology. 2023/03/01 2023;21(3):162–177. doi:10.1038/s41579-022-00841-7

8. Singh H, Dahiya N, Yadav M, Sehrawat N. Emergence of SARS-CoV-2 New Variants and Their Clinical Significance. The Canadian journal of infectious diseases & medical microbiology = Journal canadien des maladies infectieuses et de la microbiologie medicale. 2022;2022:7336309. doi:10.1155/2022/7336309

9. Gupta A, Madhavan MV, Sehgal K, et al. Extrapulmonary manifestations of COVID-19. Nature Medicine. 2020/07/01 2020;26(7):1017–1032. doi:10.1038/s41591-020-0968-3

10. Liu J, Li Y, Liu Q, et al. SARS-CoV-2 cell tropism and multiorgan infection. Cell Discovery. 2021/03/23 2021;7(1):17. doi:10.1038/s41421-021-00249-2

11. Carrau L, Frere JJ, Golynker I, et al. Delayed engagement of host defenses enables SARS-CoV-2 viremia and productive infection of distal organs in the hamster model of COVID-19. Science signaling. Jun 13 2023;16(789):eadg5470. doi:10.1126/scisignal.adg5470

12. Upadhyay V, Suryawanshi RK, Tasoff P, et al. Mild SARS-CoV-2 infection results in long-lasting microbiota instability. mBio. Aug 31 2023;14(4):e0088923. doi:10.1128/mbio.00889-23

13. Sencio V, Machelart A, Robil C, et al. Alteration of the gut microbiota following SARS-CoV-2 infection correlates with disease severity in hamsters. Gut microbes. Jan-Dec 2022;14(1):2018900. doi:10.1080/19490976.2021.2018900

14. Seibert B, Cáceres CJ, Carnaccini S, et al. Pathobiology and dysbiosis of the respiratory and intestinal microbiota in 14 months old Golden Syrian hamsters infected with SARS-CoV-2. PLoS pathogens. Oct 2022;18(10):e1010734. doi:10.1371/journal.ppat.1010734

15. Rosenke K, Meade-White K, Letko M, et al. Defining the Syrian hamster as a highly susceptible preclinical model for SARS-CoV-2 infection. Emerging microbes & infections. Dec 2020;9(1):2673–2684. doi:10.1080/22221751.2020.1858177

16. Imai M, Iwatsuki-Horimoto K, Hatta M, et al. Syrian hamsters as a small animal model for SARS-CoV-2 infection and countermeasure development. Proceedings of the National Academy of Sciences of the United States of America. Jul 14 2020;117(28):16587–16595. doi:10.1073/pnas.2009799117

17. Boudewijns R, Thibaut HJ, Kaptein SJF, et al. STAT2 signaling restricts viral dissemination but drives severe pneumonia in SARS-CoV-2 infected hamsters. Nat Commun. Nov 17 2020;11(1):5838. doi:10.1038/s41467-020-19684-y

18. Case JB, Bailey AL, Kim AS, Chen RE, Diamond MS. Growth, detection, quantification, and inactivation of SARS-CoV-2. Virology. Sep 2020;548:39–48. doi:10.1016/j.virol.2020.05.015

19. (ECDC) ECfDPaC. Threat assessment: Emergence of SARS-CoV-2 B.1.617 variants. Stockholm: ECDC. 2021 [cited 2024 Dec 22]

20. Viana R, Moyo S, Amoako DG, et al. Rapid epidemic expansion of the SARS-CoV-2 Omicron variant in southern Africa. Nature. Mar 2022;603(7902):679–686. doi:10.1038/s41586-022-04411-y

21. Chakraborty C, Bhattacharya M, Sharma AR, Dhama K, Lee SS. The rapid emergence of multiple sublineages of Omicron (B.1.1.529) variant: Dynamic profiling via molecular phylogenetics and mutational landscape studies. Journal of infection and public health. Nov 2022;15(11):1234–1258. doi:10.1016/j.jiph.2022.10.004

22. Tegally H, Moir M, Everatt J, et al. Emergence of SARS-CoV-2 Omicron lineages BA.4 and BA.5 in South Africa. Nat Med. Sep 2022;28(9):1785–1790. doi:10.1038/s41591-022-01911-2

23. Tostanoski LH, Wegmann F, Martinot AJ, et al. Ad26 vaccine protects against SARS-CoV-2 severe clinical disease in hamsters. Nature Medicine. 2020/11/01 2020;26(11):1694-1700. doi:10.1038/s41591-020-1070-6

24. Bolyen E, Rideout JR, Dillon MR, et al. Reproducible, interactive, scalable and extensible microbiome data science using QIIME 2. Nat Biotechnol. Aug 2019;37(8):852–857. doi:10.1038/s41587-019-0209-9

25. Callahan BJ, McMurdie PJ, Rosen MJ, Han AW, Johnson AJ, Holmes SP. DADA2: High-resolution sample inference from Illumina amplicon data. Nat Methods. Jul 2016;13(7):581–3. doi:10.1038/nmeth.3869

26. DeSantis TZ, Hugenholtz P, Larsen N, et al. Greengenes, a chimera-checked 16S rRNA gene database and workbench compatible with ARB. Applied and environmental microbiology. Jul 2006;72(7):5069–72. doi:10.1128/aem.03006-05

27. Faith DP. Conservation evaluation and phylogenetic diversity. Biological Conservation. 1992/01/01/ 1992;61(1):1-10. 10.1016/0006-3207(92)91201-3

28. Jaccard P. Nouvelles recherches sur la distribution florale. Bulletin de la Société vaudoise des sciences naturelles. 1908;44:223--270.

29. Baptiste Auguie AA. gridExtra: Miscellaneous Functions for "Grid" Graphics. R package version 2.3. Vienna (Austria): Comprehensive R Archive Network (CRAN). 2017 [cited 2024 Dec 22] doi:10.32614/CRAN.package.gridExtra

30. Bisanz J. qiime2R: Importing QIIME2 artifacts and associated data into R sessions. Version 0.99.6. GitHub repository. 2022 [cited 2024 Dec 22]

31. Constantin Ahlmann-Eltze IP. ggsignif: R Package for Displaying Significance Brackets for ’ggplot2. PsyArxiv (preprint). 2021;doi:doi:10.31234/osf.io/7awm6

32. Conway JR, Lex A, Gehlenborg N. UpSetR: an R package for the visualization of intersecting sets and their properties. Bioinformatics. Sep 15 2017;33(18):2938–2940. doi:10.1093/bioinformatics/btx364

33. Kassambara A. ggpubr: ’ggplot2’ Based Publication Ready Plots. R package version 0.7.2. Vienna (Austria): Comprehensive R Archive Network (CRAN). 2023 [cited 2024 Dec 22] doi:10.32614/CRAN.package.ggpubr

34. Kassambara A. rstatix: Pipe-Friendly Framework for Basic Statistical Tests. R package version 0.7.0. Vienna (Austria): Comprehensive R Archive Network (CRAN) 2023 [cited 2024 Dec 22] doi:10.32614/CRAN.package.rstatix

35. Neuwirth E. RColorBrewer: ColorBrewer Palettes. R package version 1.1-3. Vienna (Austria): Comprehensive R Archive Network (CRAN). 2022 [cited 2024 Dec 22] doi:10.32614/CRAN.package.RColorBrewer

36. Uhlén M, Fagerberg L, Hallström BM, et al. Tissue-based map of the human proteome. 2015;347(6220):1260419. doi:doi:10.1126/science.1260419

37. Jung K, Wang Q, Scheuer KA, Lu Z, Zhang Y, Saif LJ. Pathology of US porcine epidemic diarrhea virus strain PC21A in gnotobiotic pigs. Emerging infectious diseases. Apr 2014;20(4):662–5. doi:10.3201/eid2004.131685

38. Li X, Song Y, Wong G, Cui J. Bat origin of a new human coronavirus: there and back again. Science China Life sciences. Mar 2020;63(3):461–462. doi:10.1007/s11427-020-1645-7

39. Stein SR, Ramelli SC, Grazioli A, et al. SARS-CoV-2 infection and persistence in the human body and brain at autopsy. Nature. Dec 2022;612(7941):758–763. doi:10.1038/s41586-022-05542-y

40. Xiao F, Tang M, Zheng X, Liu Y, Li X, Shan H. Evidence for Gastrointestinal Infection of SARS-CoV-2. Gastroenterology. May 2020;158(6):1831–1833.e3. doi:10.1053/j.gastro.2020.02.055

41. Zhou J, Li C, Liu X, et al. Infection of bat and human intestinal organoids by SARS-CoV-2. Nat Med. Jul 2020;26(7):1077–1083. doi:10.1038/s41591-020-0912-6

42. Marasco G, Cremon C, Barbaro MR, et al. Prevalence of Gastrointestinal Symptoms in Severe Acute Respiratory Syndrome Coronavirus 2 Infection: Results of the Prospective Controlled Multinational GI-COVID-19 Study. The American journal of gastroenterology. Jan 1 2022;117(1):147–157. doi:10.14309/ajg.0000000000001541

43. Whitaker M, Elliott J, Bodinier B, et al. Variant-specific symptoms of COVID-19 in a study of 1,542,510 adults in England. Nat Commun. Nov 11 2022;13(1):6856. doi:10.1038/s41467-022-34244-2

44. Mautner L, Hoyos M, Dangel A, Berger C, Ehrhardt A, Baiker A. Replication kinetics and infectivity of SARS-CoV-2 variants of concern in common cell culture models. Virol J. Apr 26 2022;19(1):76. doi:10.1186/s12985-022-01802-5

45. Toomer G, Burns W, Garcia L, et al. Characterization of Three Variants of SARS-CoV-2 In Vivo Shows Host-Dependent Pathogenicity in Hamsters, While Not in K18-hACE2 Mice. Viruses. Nov 21 2022;14(11)doi:10.3390/v14112584

46. Yuan S, Ye ZW, Liang R, et al. Pathogenicity, transmissibility, and fitness of SARS-CoV-2 Omicron in Syrian hamsters. Science. Jul 22 2022;377(6604):428–433. doi:10.1126/science.abn8939

47. Suzuki R, Yamasoba D, Kimura I, et al. Attenuated fusogenicity and pathogenicity of SARS-CoV-2 Omicron variant. Nature. Mar 2022;603(7902):700–705. doi:10.1038/s41586-022-04462-1

48. Mohandas S, Yadav PD, Sapkal G, et al. Pathogenicity of SARS-CoV-2 Omicron (R346K) variant in Syrian hamsters and its cross-neutralization with different variants of concern. EBioMedicine. May 2022;79:103997. doi:10.1016/j.ebiom.2022.103997

49. Suresh V, Parida D, Minz AP, Sethi M, Sahoo BS, Senapati S. Tissue Distribution of ACE2 Protein in Syrian Golden Hamster (Mesocricetus auratus) and Its Possible Implications in SARS-CoV-2 Related Studies. Frontiers in pharmacology. 2020;11:579330. doi:10.3389/fphar.2020.579330

50. Lehmann M, Allers K, Heldt C, et al. Human small intestinal infection by SARS-CoV-2 is characterized by a mucosal infiltration with activated CD8(+) T cells. Mucosal immunology. Nov 2021;14(6):1381–1392. doi:10.1038/s41385-021-00437-z

51. Tian Y, Rong L, Nian W, He Y. Review article: gastrointestinal features in COVID-19 and the possibility of faecal transmission. Alimentary pharmacology & therapeutics. May 2020;51(9):843–851. doi:10.1111/apt.15731

52. Termansen MB, Frische S. Fecal-oral transmission of SARS-CoV-2: A systematic review of evidence from epidemiological and experimental studies. American journal of infection control. Dec 2023;51(12):1430–1437. doi:10.1016/j.ajic.2023.04.170

53. Joshi M, Mohandas S, Prasad S, et al. Lack of evidence of viability and infectivity of SARS-CoV-2 in the fecal specimens of COVID-19 patients. Frontiers in public health. 2022;10:1030249. doi:10.3389/fpubh.2022.1030249

54. Wurtzer S, Lacote S, Murri S, et al. Reduction in SARS-CoV-2 Virus Infectivity in Human and Hamster Feces. Viruses. Aug 15 2022;14(8)doi:10.3390/v14081777

55. Bernard-Raichon L, Venzon M, Klein J, et al. Gut microbiome dysbiosis in antibiotic-treated COVID-19 patients is associated with microbial translocation and bacteremia. Nat Commun. Nov 1 2022;13(1):5926. doi:10.1038/s41467-022-33395-6

56. Yeoh YK, Zuo T, Lui GC, et al. Gut microbiota composition reflects disease severity and dysfunctional immune responses in patients with COVID-19. Gut. Apr 2021;70(4):698–706. doi:10.1136/gutjnl-2020-323020

57. Arjmand B, Ghorbani F, Koushki M, Rezai-Tavirani M. Gastrointestinal symptoms in patients with mild and severe COVID-19: a scoping review and meta-analysis. Gastroenterology and hepatology from bed to bench. Fall 2020;13(4):321–330.

