## Supplementary material for "SARS-CoV-2 variants exhibit differential gut tropism and dysbiosis in Syrian golden hamsters": Suuplemetary material

**Affiliation:**

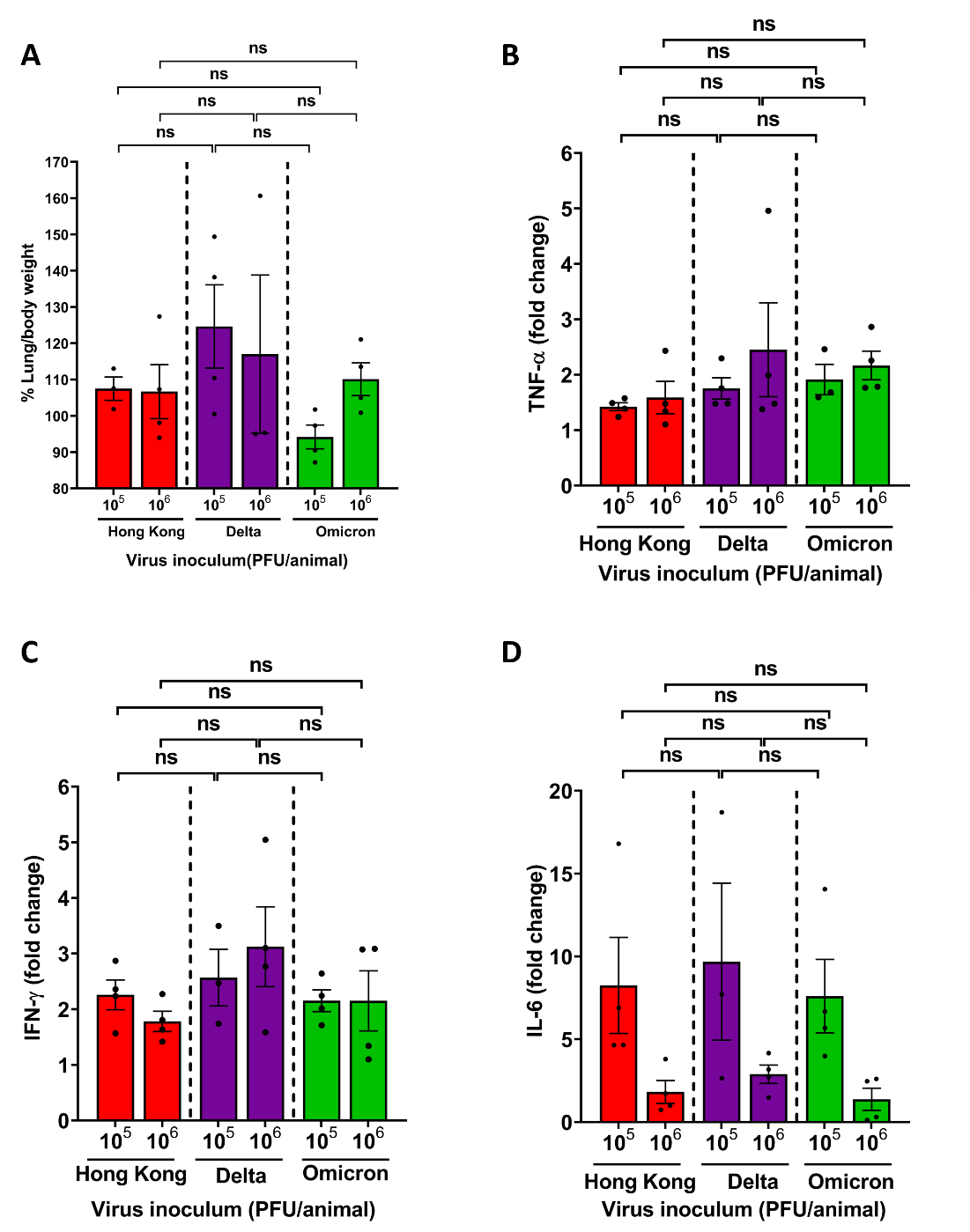

**Supplemental Figure 1. Tissue inflammation and inflammatory cytokine levels in the hamster lungs infected with SARS-CoV-2 VOCs.** Hamsters (n=3 to 4) were infected with either 10^5^ or 10^6^ PFU/animal of SARS-CoV-2 Hong Kong strain, or VOCs Delta or Omicron. **(A)** Animals were sacrificed at 4 dpi, lung weight was measured and lung/body weight ratio was plotted. The total RNA was extracted from the lung and used to quantify expression levels of **(B)** TNF-α, **(C)** IFN-ϒ, and **(D)** IL-6 by RT-qPCR. Statistics were performed by one-way ANOVA with Tukey's multiple comparison test. ns - not significant. Error bars indicate Mean ± SEM.

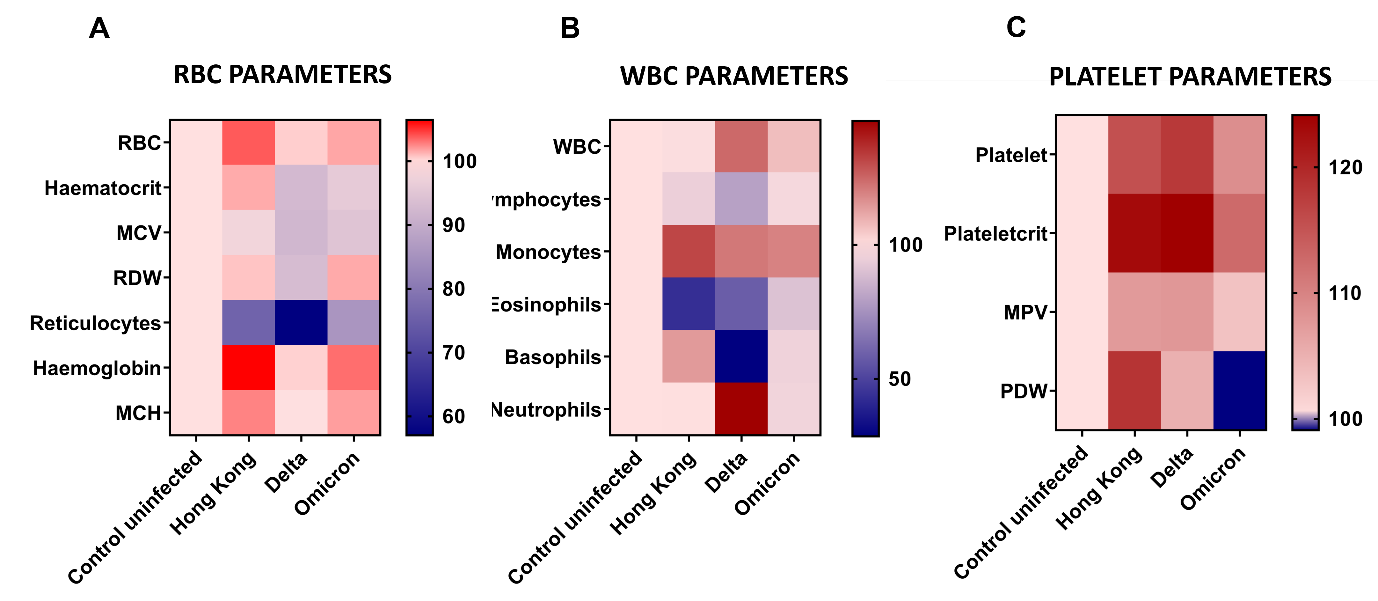

**Supplemental Figure 2. Hamsters infected with the Delta variant showed the highest altered haematological parameters.** Hamsters (n=3 to 4) were infected with 10^6^ PFU/animal of SARS-CoV-2 Hong Kong, Delta, or Omicron, and blood was collected on day 4 post-infection. Different haematological parameters of **(A)** RBC, **(B)** WBC, and **(C)** platelet levels were measured.

**Table S1: Sequences of primer and probes used for RT-qPCR**

| **Gene** | **Primer sequence ( 5’-3’)** |
| --- | --- |
| **Nucleocapsid-1** | **Forward:** GACCCCAAAATCAGCGAAAT  **Reverse:** TCTGGTTACTGCCAGTTGAATCTG |
| **IFN-γ** | **Forward**: TGTTGCTCTGCCTCACTCAGG3  **Reverse:** AAGACGAGGTCCCCTCCATTC3 |
| **IL-6** | **Forward:** AGGCCATCCTGATGGAGAAG  **Reverse:** GGTATGCTAAGGCACAGCAC |
| **TNF-α** | **Forward:** TGAGCCATCGTGCCAATG  **Reverse:** AGCCCGTCTGCTGGTATC |
|  | **Probe Sequence ( 5’-3’)** |
| **Nucleocapsid-1** | (6-FAM/BHQ-1) ACCCCGCATTACGTTTGGTGGACC |

**Figure S1. Tissue inflammation and inflammatory cytokine levels in the hamster lungs infected with SARS-CoV-2 VOCs**

**Figure S2. Hamsters infected with the Delta variant showed the highest altered haematological parameters.**

**Table S1: Sequences of primer and probes used for RT-qPCR**
